# InterPred: A pipeline to identify and model protein-protein interactions

**DOI:** 10.1101/080754

**Authors:** Claudio Mirabello, Björn Wallner

## Abstract

Protein-protein interactions (PPI) are crucial for protein function. There exist many techniques to identify PPIs experimentally, but to determine the interactions in molecular detail is still difficult and very time-consuming. The fact that the number of PPIs is vastly larger than the number of individual proteins makes it practically impossible to characterize all interactions experimentally. Computational approaches that can bridge this gap and predict PPIs and model the interactions in molecular detail are greatly needed. Here we present InterPred, a fully automated pipeline that predicts and model PPIs from sequence using structural modelling combined with massive structural comparisons and molecular docking. A key component of the method is the use of a novel random forest classifier that integrate several structural features to distinguish correct from incorrect protein-protein interaction models. We show that InterPred represents a major improvement in protein-protein interaction detection with a performance comparable or better than experimental high-throughput techniques. We also show that our full-atom protein-protein complex modelling pipeline performs better than state of the art protein docking methods on a standard benchmark set. In addition, InterPred was also one of the top predictors in the latest CAPRI37 experiment.

InterPred source code can be downloaded from http://wallnerlab.org/InterPred

## 1 Introduction

Protein-protein interactions (**PPI**) are crucial for many cellular functions, such as signal transduction, transport, metabolism, and transcription. Since knowledge of PPIs is important to understand both basic biology and human disease at the molecular level [1], major efforts have been devoted to experimentally characterize PPIs from the level of detecting interactions to exact molecular details of the interaction [2].

Detection and identification of protein interactions can be experimentally performed using high-throughput (HT) techniques such as yeast two-hybrid [3] and affinity purification [4]. But these methods have inherent limitations resulting in many false positives and negatives [5]. Novel methods using proximity-ligation techniques like BioID [6] are promising, but are limited to finding proteins that are part of the same complex and not necessarily in direct physical interaction. Characterization of the quaternary structure of proteins in molecular detail can still only be performed for individual protein complexes, using regular structural biology methods like x-ray crystallography, NMR or cryo-EM.

Complementing the experimental methods, several computational approaches have also been derived to predicted if proteins interact from the amino acid sequence [7, 8, 9], using co-evolution [10], gene co-expression [11], and phylogenetic profiles [12] or by combining different sources of information [13].

Similarly, there are many methods for predicting the quaternary structures of proteins, however still mostly restricted to dimers. These methods can be categorized as template-based or template-free methods. Template-based modelling methods identify complex structure templates by aligning the amino acid sequences [14, 15, 16] or structural models [17, 18] of the target chains against solved complex structures in the PDB (Protein Data Bank) or libraries of the complex interface [19, 20, 21, 22]. In template-free methods complex structures are constructed by assembling known or modelled monomers using protein-protein docking [23, 24, 25, 26].

Both categories of methods have their advantages and disadvantages. Template-free modelling methods have the advantage that they can be applied to any protein pair given that the monomer structures are known or can be modelled. But in general, the quality of the prediction cannot be guaranteed in particular if the monomer structures change upon binding [27]. A further limitation is that many thousands of docking models must be sampled to find at least one correct docking model [28], making them unsuitable for genome-wide applications.

Template-based methods have in general higher accuracy if there are homologous complex templates available. The accuracy depends heavily on the evolutionary distance between the target and template, and drops rapidly when the sequence identity approaches the twilight-zone (i.e. when the sequence identity is around 30%). However, it has been shown that protein-protein interfaces are degenerate and that in distantly and even unrelated proteins often are similar [29, 30, 31]. This means that template-based modelling could potentially be expanded using analogous protein complex template structures.

Since structure is more conserved than sequence, methods that compare structural models, as opposed to sequences, against a template library should have a higher chance of finding analogous templates. Indeed, by conducting the structural search more broadly to find similar interfaces from proteins not necessarily homologous, combined with features extracted from structural alignments with non-structural clues in a Bayesian classifier has enabled the identification of PPIs on a genome-wide scale (human, yeast) with accuracy that is comparable to HT experimental techniques [32]. Other methods have also reported improved performance in docking model quality using structural alignment searches [18, 22]. However, despite the seemingly large number of available methods for template-based docking using structural alignments [18, 22, 33, 34, 35], none of them, to the best of our knowledge, is available for practical large-scale use outside the lab they were developed.

In this study, we present InterPred, a computational pipeline to predict and model protein-protein interactions using structural modelling combined with massive structural comparisons and molecular docking. A key component of the method is the use of a random forest classifier to integrate several structural features to distinguish correct from incorrect protein interaction models. The method is open source and available as a stand-alone download.

## 2 Methods

The aim of InterPred is two-fold: 1) to decide if two proteins interact and 2) If so, predict the molecular details of the interaction.

In short, the InterPred pipeline consists of three steps (Figure 1): (1) target homology modelling, (2) template search including interaction modelling/scoring, and (3) refinement.

In the first step, the sequences of the protein targets, whose interaction is investigated, are used to build structural models using homology modelling. Next, structural alignments are used to find close and distant structural similarities to the two models in the Protein Data Bank (PDB) [36]. Whenever the structures similar to the two models also form a complex in the PDB, it defines an interaction template for modelling the interaction. An interaction model is then built by superimposing the representative structures to their corresponding structural homologs in the interaction template. Based on the interaction model a set of features are calculated and used as input to a random forest classifier trained to sift out the more promising models that will go through to the final refinement step. Each step is described in detail below.

**Figure 1:**
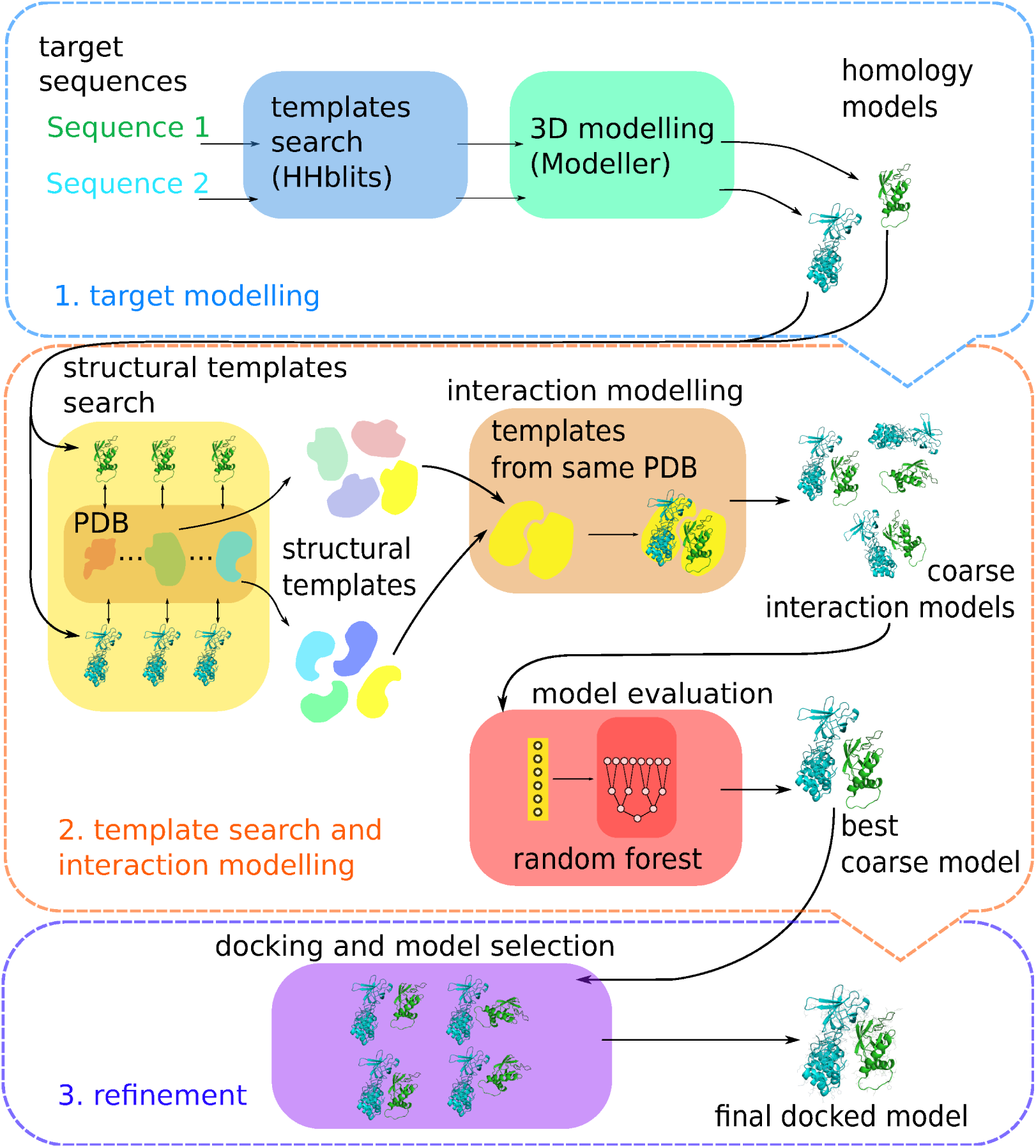
Overview of the InterPred pipeline

### 2.1 Structural Modelling of Target Sequences

The construction of structural models for the target sequences is a crucial step towards the search for structural templates, since without any models it is not possible to run InterPred at all. The target sequences are used in a homology modelling system combining HHblits [37] from the HHpred suite for the template search and MODELLERv9.13 [38] to build full-atom 3D models. The Hidden Markov Model (HMM) profiles are built by searching the *uniprot20_2013_03* database bundled in the HHpred suite, using two iterations of HHblits with Expect value (E-value) cut-off 10^−2^ and maximum pairwise identity 90%. The HMM profile was then used to search against the HHpred PDB database clustered at 70% sequence identity (September 6, 2014) with the same settings. Models were constructed using the template with the lowest E-value and for each non-overlapping template with E<10^−3^ using MODELLER. If different templates cover different regions of the target sequence, models are built separately for each region. This is particular useful when building models of multi-domain proteins where no homology model can be found spanning across all domains. This way, additional domains that would otherwise be overlooked will still be used in the template search step. Moreover, it is preferable to run the structural template search with single domain structures rather than a large multi-domain structure. Because the size difference between two domains during the structural search could cause a bias towards templates that are similar to the larger domain and potentially useful templates for the smaller domain could be missed.

### 2.2 Template Search and Interaction Modelling

To model the interaction, potential interaction templates are identified by first defining the space of similar structures for each of the structures from the target modelling step, and then by searching for overlaps between spaces, i.e. cases were the same experimental structure occurs in both spaces.

The space of similar structures is defined as structures with maximum TM-score>0.5 (normalized by the shortest length) using structural alignments with TM-align [39], against every chain in the PDB using both biological and asymmetric unit. In addition, to avoid random hits occupying the spaces we employ a threshold on minimum length of the structural alignment (L>10). It was observed that some (5%) alignments passing these thresholds still had a large Root Mean Square Deviation (RMSD>4Å) of the aligned positions. Since it has been shown that docking success rate drops significantly for models with RMSD>4Å [40], we decided to also filter these alignments out. positions.

For each potential interaction template, a coarse interaction model is obtained in the *interaction modelling* step, by superimposing the structural representatives of the targets onto the interaction template, thereby transferring the positional relationship of the template to the targets. However, it is important to note that not all of the interaction models constructed in this way will be correct. And given multiple interaction models for the same targets, a ranking method is needed to select those that are potentially better given the features at hand.

### 2.3 Ranking Interaction Models

To construct a ranking of plausible interaction models, a random forest classifier was trained to predict the likelihood of interaction based on the series of features (Table 1). All features are only based on properties that can be calculated from the model itself. There is other useful information that could potentially improve performance, co-expression levels, gene ontology similarity, functional similarity and phylogenetic profiles, or whether the two proteins are essential for survival [13]. However, none of these features will be specific to the 3D coordinates of a particular interaction model making them unsuitable for ranking interaction models. Another problem with using these type features is how to deal with cases when information is missing, which is the most likely scenario. Thus, to avoid these problems we chose to train the random forest classifier only on structural information at this stage. The structural features are described in detail below.

**Table 1.**
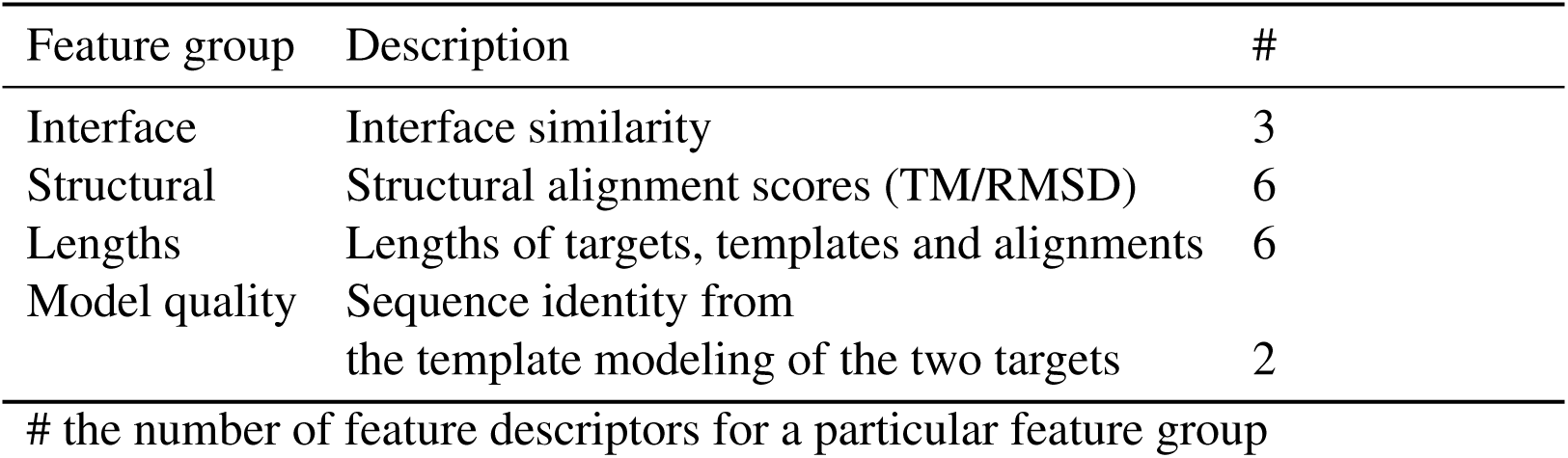
Description of features used to train the random forest classifier.

| Feature group | Description | # |
| --- | --- | --- |
| Interface | Interface similarity | 3 |
| Structural | Structural alignment scores (TM/RMSD) | 6 |
| Lengths | Lengths of targets, templates and alignments | 6 |
| Model quality | Sequence identity from<br>the template modeling of the two targets | 2 |
| # the number of feature descriptors for a particular feature group |  |  |

*Interface features* describe the similarity between the interface of the template and the interface of the model. The interface between two molecules is all residue pairs from each molecule in the complex, for which any heavy atom is within a certain distance cutoff. Common cutoffs found in the literature are the sum of the van der Waals radii of the atoms plus a threshold of 0.5Å [41], 4.5 Å [19], 5.0Å [42]. While this works well when dealing with native structures, the interacting chains in coarse models can sometimes be placed further apart than those in the corresponding template. Using such stringent thresholds will, then, often cause the interface in the coarse model to be “empty”, i.e. no residues between the two chains are within the cutoff. To avoid this, we empirically increment the cutoff to 6.5A.

The interface features are described by the size of the interface in the template complex, model complex, and the number of residues that overlaps when the model is superimposed on the template (Figure 2). The number overlapping residues are calculated similarly to how it was done in [40], although as an absolute number rather than normalized by the number of total interacting residues.

**Figure 2.**
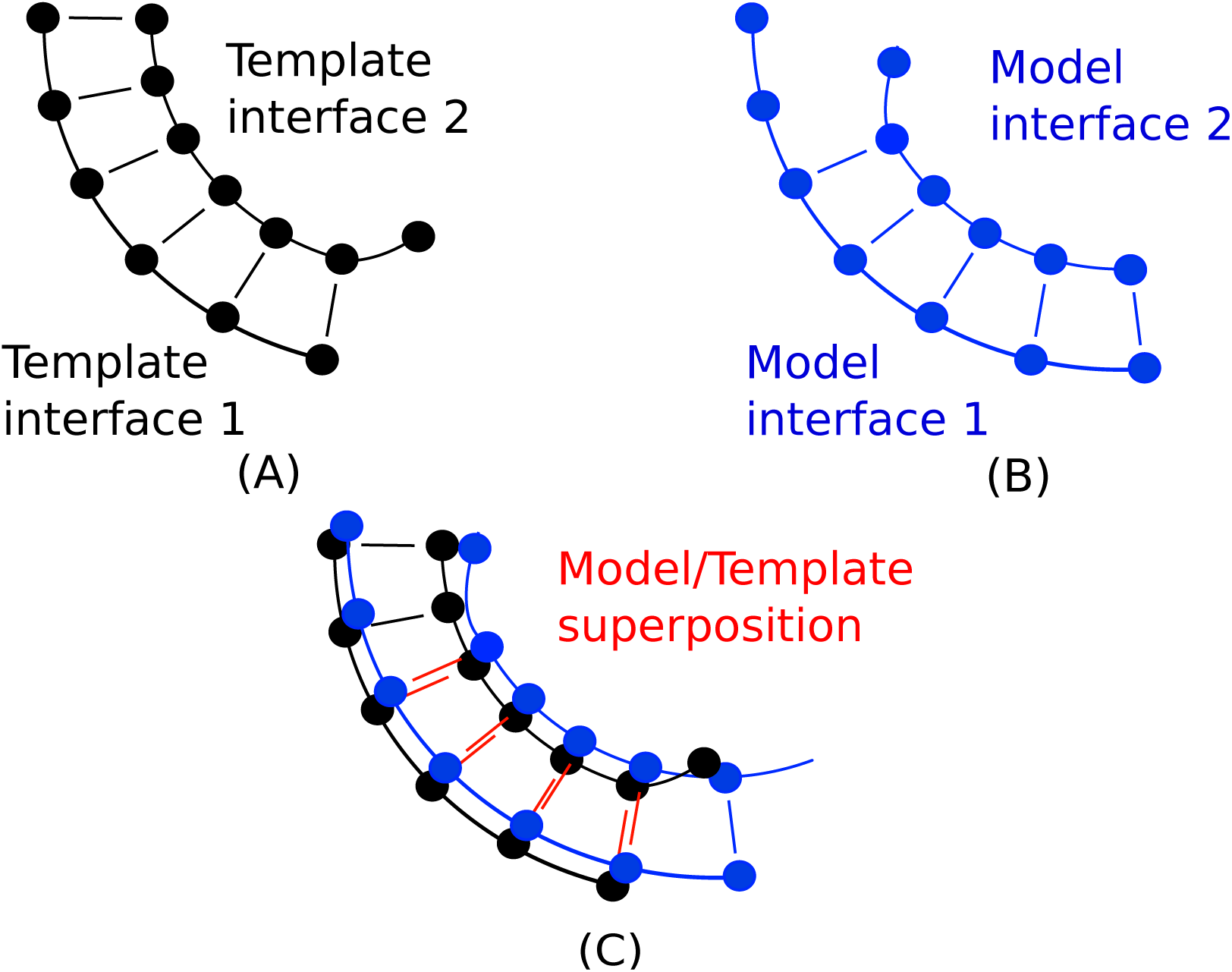
The interface features describe the similarity between the interface of the template and the model by three numbers: the number of residues in the template complex interface (solid lines in A), the number of residues model complex interface (solid lines in B), and the number of residues that are shared when the template and model are superimposed (double lines in C).

*Structural alignment features* describe the quality of the structural alignment between the two target structures and their structural templates using the RMSD for the superposition, TM-score normalized by the length of target, and TM-score normalized by the length of the template, for the two targets separately.

*Model quality features* describe the quality of the sequence alignment used to build the target structures from the sequence input and was represented by the sequence identity between the sequence templates and the target sequences for the two models.

We also tried using the sequence similarity of the target-template structural alignment as a feature, but it did not improve performance (data not shown)

### 2.4 Interaction Model Refinement

As the interaction models are based purely on superpositions, they might contain severe clashes and sub optimal interactions. To produce all-atom models with no clashes and optimized interactions, interaction models were refined using RosettaDock [43] with the perturbation flag (*–dock_pert)* set to “5 12”, for the standard deviations in Angstrom and degrees for the initial translation and rotation perturbation, respectively. 10,000 decoys were generated and ranked using interface RMSD (IRMSD) to the starting conformation, i.e. the decoys that changed their interface least were selected.

### 2.5 Data sets

For training and testing various aspects of InterPred four different protein-protein interaction sets were used. A binary protein-protein interaction set (1), a binary interaction model set (2), an interaction set from a recent benchmark of protein-protein interaction experiments (3), and finally a 3D interaction set was used to assess the refinement and the ability to predict the 3D structure of the interaction models (4).

#### 2.5.1 Binary protein-protein interaction set

The binary protein-protein interaction set consist of positive and negative interactions and was constructed in a similar way to [13]: the set of interacting pairs, or *positive set,* is composed of yeast and human protein pairs that have been shown to interact by at least two separate publications. The set of non-interacting pairs, or *negative set,* consists of human and yeast protein pairs from different cellular compartments according to Gene Ontology [44]. To make absolutely sure that none of the pairs are interacting, we were strict in picking proteins that are located in one and only one of the following compartments: membrane, mitochondria, endoplasmic reticulum, and nucleoplasm. The resulting positive set contained 30,247 interacting pairs and the negative set contained 13,121 non-interacting pairs.

#### 2.5.2 Binary interaction model set

To train the random forest classifier, a binary interaction model set consisting of correct and incorrect structural interaction models were constructed by running the first (homology modelling) and second (template search and interaction modelling) steps of the pipeline on the binary protein-protein interaction set, generating on average one to two thousand interaction models per protein pair, in total 64-million models for the positive and 14-million models for the negative pairs. The interaction models based on the negative sets are all incorrect and can thus all be used as negative interaction models. However, for the interaction models based on the positive set it cannot be assumed that all 64-million possible interaction models are correct. In fact, most of them will be incorrect, since the total number of possible interaction is much higher that the subset of correct interactions. To make sure that positive interaction models are correct only the 4,162 protein pairs for which obvious homologous interaction templates could be found using sequences alone were included in the *correct interaction model set.* This was done by searching for sequence homologs in the PDB for each protein in the positive protein-protein interaction set using HHblits and construct interaction models for those pairs which shared sequence templates from the same PDB. The final set used for training consisted of 14-million incorrect and 80,921 correct interaction models.

### 2.6 Cross-validation

To prepare the sets for 10-fold cross-validation during random forest training, the full set of sequences from binary protein-protein interaction set were clustered using BLASTCLUST [45] and divided into 10 parts in such a way that no pair of targets from two different folds shared more than 50% sequence similarity at 90% coverage. The exact BLASTCLUST parameters are not crucial, since the training is not performed directly on sequences, but on a rather limited set of features calculated from the interaction models. The aim of the clustering was to ensure that no two sets of features from two different folds were identical.

#### 2.6.1 Braun benchmark set

The performance of InterPred to predict protein-protein interactions was compared to several high-throughput methods on a benchmark set developed by Braun et *al.* (Braun benchmark set) consisting of 92 interacting and 92 non-interacting protein pairs [46].

#### 2.6.2 BM4: Docking model quality benchmark

To test the capability of InterPred to produce and dock interaction models, a set of protein-protein complexes from the docking Benchmark 4.0 [47] was used (BM4). This benchmark consists of 176 bound targets and the corresponding unbound interactors.

### 2.7 Random forest classifier

The random forest classifier in the TreeBagger class from Matlab’s Statistics and Machine Learning Toolbox (version R2014b) were used. An ensemble of 100 decision trees per forest was trained to recognize correct and incorrect interactions. The fraction of decision trees predicting a positive interaction determines the *InterPred score.* To find the best combination of features, several versions of the classifier were trained using different combinations of input features. The trainings were performed on the *binary interaction model set* using 10-fold crossvalidation as described above. The final testing was performed on protein pairs from the full binary protein-protein interaction set using the same cross-validation sets. The InterPred score for a protein-protein pair was the highest InterPred score for any interaction model generated from that pair.

### 2.8 Performance measures

To measure the detection performance, receiver operating characteristic (ROC) curves showing True Positive Rate (TPR) against False Positive Rate (FPR), were used

FPR is defined as:

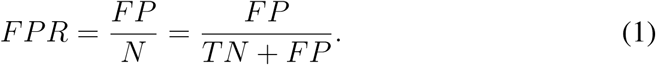

TPR is defined as:

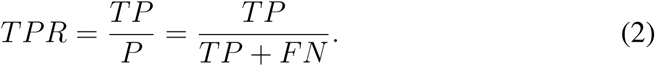
where P is the number of positive examples in the set (e.g. number of interacting pairs), *N* the number of negative examples, *TP* the number of correctly identified positives and FP the number of incorrectly identified positives.

The Likelihood Ratio (LR) is then defined as:

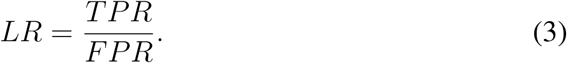

where TPR and *FPR* are defined above.

#### 2.8.1 IS-score

To assess the quality of refined and coarse interaction models in the docking quality benchmark, each model is compared to its corresponding native structure using the Interface Similarity score (IS-score) [48], which outputs a score from 0 to 1, where 0 represents a completely incorrect model and 1 a perfect model. Two thresholds (0.12 and 0.17) are applied to IS-score to distinguish between “Bad”, “Acceptable” and “Near-native” models, as done previously for the development of the PRISM method [34]. It is important to note that this classification differs from the one used in CAPRI [42], but it has been shown that IS-score correlates well with the measures that are used to calculate the CAPRI classification [34].

### 2.9 PRISM

PRISM is a widely used method based on structural templates [22] available through a web interface. The structural template search on PRISM is performed with rigid-body structural comparisons of target proteins to known template protein-protein interfaces, then a set of thresholds is employed to sift out the most promising interfaces before the refinement, based both on structural similarity and evolutionary conservation of putative binding residues step [49]. Contrarily to the approach used in InterPred, no machine learning techniques are used to rank the templates.

For comparisons to PRISM, BM4 targets were submitted to the PRISM server, and the refined models were downloaded and ranked according to the Fiberdock energy scores reported on the results page.

### 2.10 ZDOCK

ZDOCK [50] is a template-free method for docking two partner proteins using fast fourier transform correlation to exhaustively explore the translation degree of freedom, limiting the search to the rotational degrees of freedom between the two partners. ZDOCK3.0.2 was shown to be the best among 18 template-free methods in a recent benchmark using the BM4 set [28].

For comparisons to ZDOCK3.0.2, a set of interaction decoys generated by ZDOCK3.0.2 for the targets in the BM4 set were downloaded from the ZDOCK webpage [51]. This decoy set contains 3,600 ranked predictions covering the rotational degree of freedom in all possible relative docking orientation at 15 degree resolution for each test case.

## 3 Results and Discussion

In this work, we have developed InterPred, a method that predicts if two proteins interact and if so also produce a three-dimensional interaction model. The exact details of the different parts of InterPred are described in Methods. Here, we give the main results and provide a benchmark to existing methods, both in terms of the ability to detect protein-protein interactions and to model the interaction in molecular detail.

### 3.1 Detection: Training of the Random Forest

The random forest classifier was trained on the binary interaction model set (see Methods). Features were calculated from the interaction models and several random forests were trained using different combinations of features. The impact of the different features used in the training on the ability to correctly identify interacting protein-protein pairs in the full binary protein-protein interaction set was measured using a Receiver Operating Characteristic (ROC) curve (Figure 3), where the FPR ( Eq. 1) is plotted against the TPR ( Eq. 2) by varying the cutoff on the InterPred score. As more features are considered, the area under the ROC curves increases, the increase is most prominent in the low FPR region (inset in Figure 3). This is important since the number of non-interacting pairs will be several orders of magnitude larger than the number of interacting pairs. Thus, it is then vital that a system designed to detect and predict interactions will can do so at very low FPRs. Clearly, at FPR<0.005 the additional features enables detection of 50%-70% more true interactions compared to a classifier only using TM-score, e.g. 4,537 predicted true positives vs. 7,561 at FPR=0.0025.

**Figure 3:**
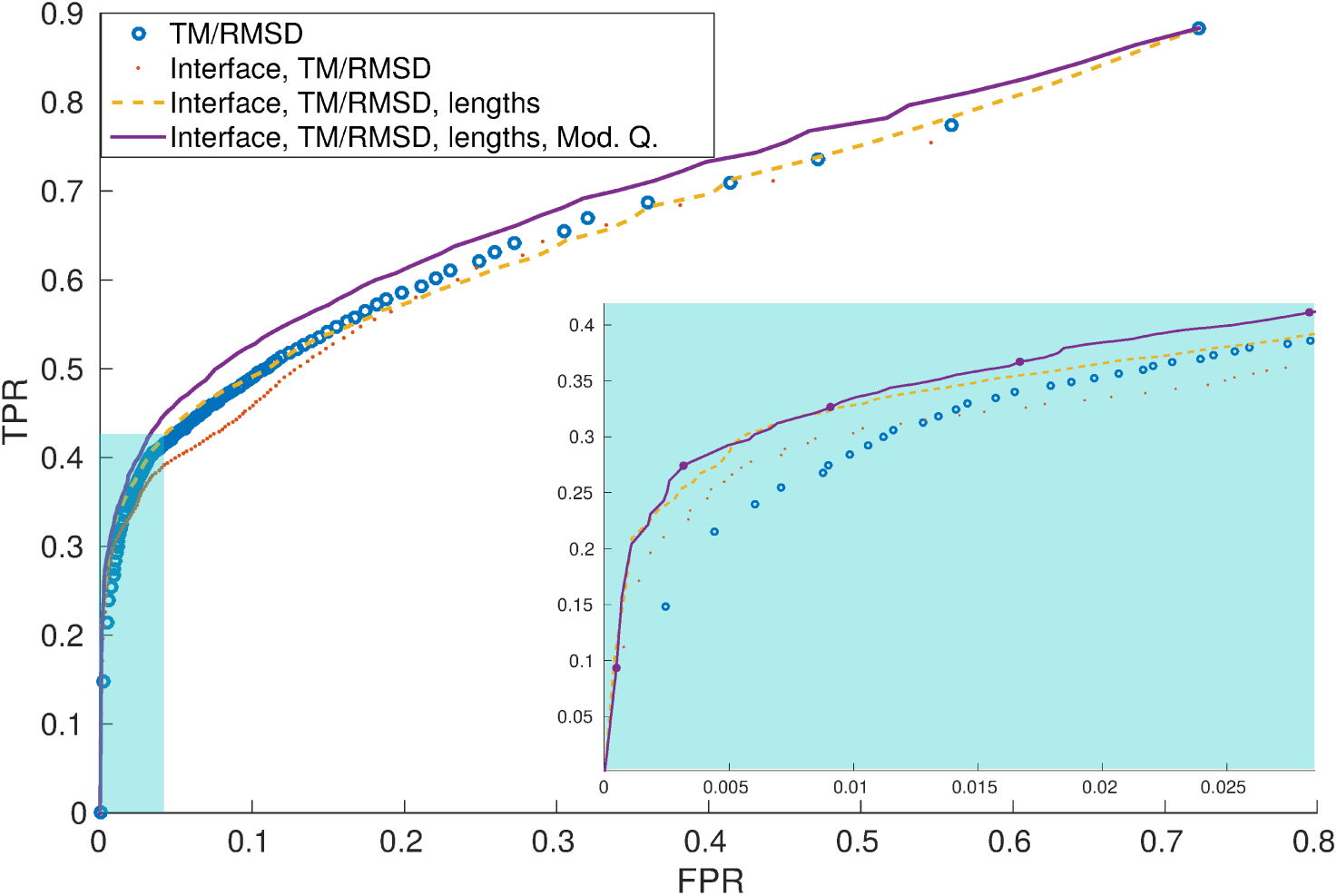
Performance for random forest classifiers trained different features. In the low FPR area (cyan), the filled circles on the curve for the final predictor highlight the thresholds on the InterPred score from 1 to 0.6 in 0.1 steps.

### 3.2 Detection: Homology modelling

The first step in the InterPred pipeline is to use homology modelling to build 3D structures from the input sequences (see Methods). It is important to model as much as possible of the input sequence, since the pipeline is based on structural comparisons. This step also models additional domains that were not covered by the best template into separate structures. No attempt is made to model the complete multi-domain protein, instead a model built from each template is used as a separate input to the pipeline.

The binary protein-protein interaction set of 43,368 target pairs contained 24,237 different proteins. For 23,397 of these at least one domain could be modelled. Modelling additional domains not covered by the best template an additional 7,960 (34%) domains could be modelled. These additional domains effectively almost double the number of interactions models, indicating the importance of model as much as possible of the target sequence.

It is well known that the quality of homology models depends to high degree on the sequence similarity to known structural templates. We included the sequence identity between the two target sequences and their respective templates at the homology modelling stage among the features for the prediction. The ROC curve in Figure 3 shows that the predictions are more accurate if this features are included, especially at low FPRs.

To assess how homology model quality influence the ability to detect correct protein-protein interactions, the binary protein-protein interaction set is split into three subsets, depending on the sequence similarity between the targets and templates used to build the models: hard (<20%), medium (20%-40%), and easy (> 40%). We calculated the Likelihood Ratio (LR, see Methods) for different InterPred score thresholds over the whole set and for each of the three subsets (Figure 4). Overall, the LR for the hard cases are not so much different from the LR for the easy cases. Indicating that there is still possible to make good predictions even using models based on remote homologs. In fact, for the medium cases there are no false positives for InterPred score >0.9. Also, the fact that we are using the model quality feature in the random forest predictor might help balancing the performance over the three sets.

**Figure 4.**
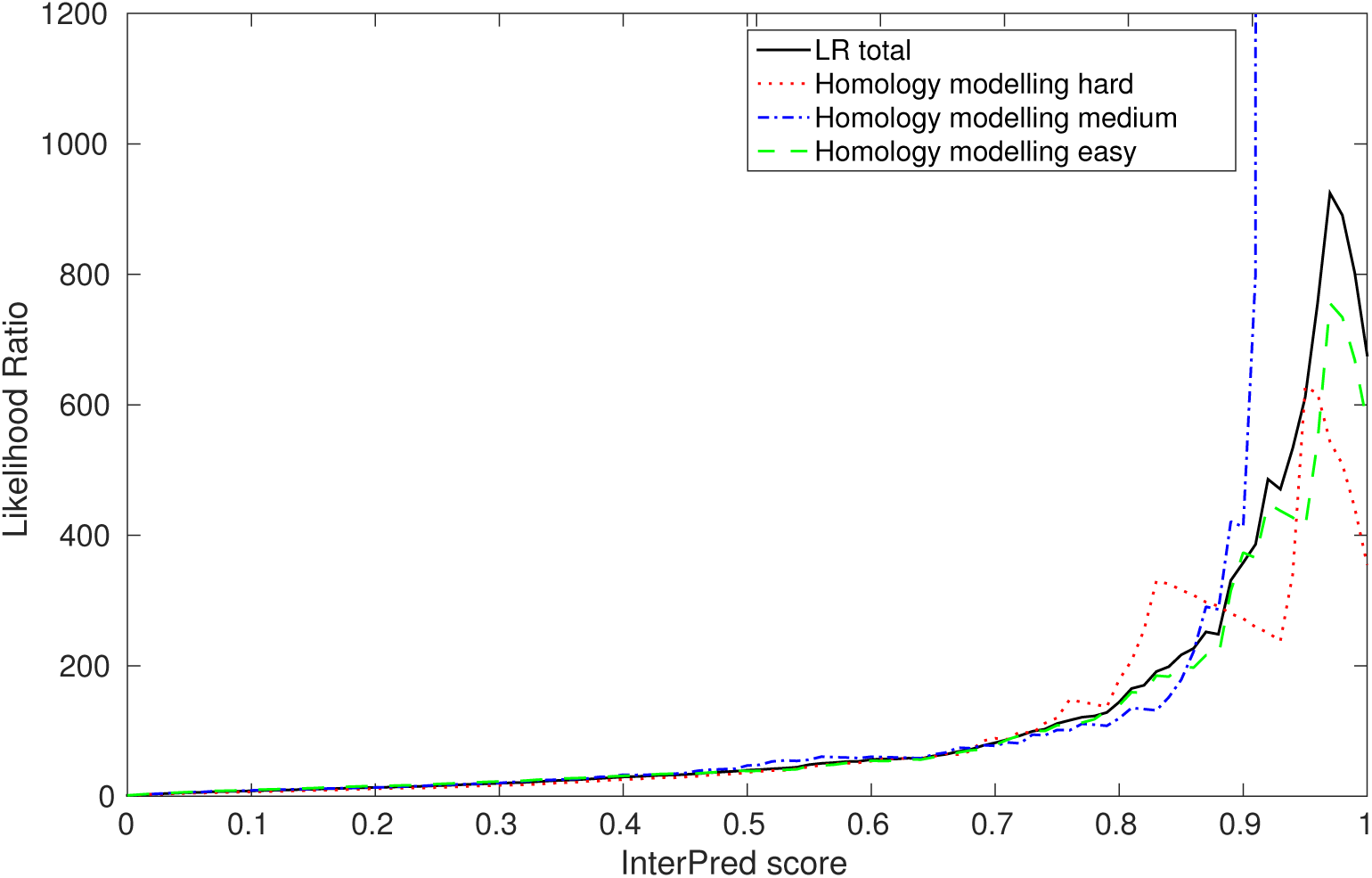
Effects of the quality of the homology models during the detection test: the Likelihood Ratio (LR) against the InterPred score for all, hard, medium, and easy homology modelling cases.

### 3.3 Detection: Braun benchmark results

We compare InterPred to a battery of four complementary, high-throughput protein interaction assays: Yeast two-hybrid (Y2H) [52], mammalian protein-protein interaction trap (MAPPIT) [53], luminescence-based mammalian interactome (LUMIER) [54], protein complementation assay (PCA) [55] and a modified version of the nucleic acid programmable protein array (wNAPPA) [56]. The comparison was performed on the Braun benchmark set [46] (see Methods).

To ensure a fair comparison, we removed any pair from the training set of InterPred where both partners have sequence identity ≥ 50% to any target pair in the Braun benchmark set and retrained InterPred. To avoid self-hits, e.g. target pairs for which the complex structure has already been experimentally resolved, we removed any structure in the PDB library that has sequence identity ≥ 90% to any target in the benchmark.

The resulting ROC curves (Figure 5), show that InterPred performs considerably better than any single high-throughput method at all FPR ranges. In particular, it correctly predicts the interaction for 25% more targets than the best high-throughput method with no false positives (39 of 92 interacting couples detected at FPR=0). We also report the performance of a consensus approach that combines all the previous experimental assays (pink circle in Figure 5), where a maximum of 55 out of the 92 interactions from the positive set could be detected at <10% FPR [46]. Our results show that InterPred performs at the same level of the consensus approach, correctly detecting the same amount of interactions at the same FPR. It is also important to know that the combined result of the experimental assays has been obtained by picking thresholds for each method so that the highest TPR could be obtained while keeping a low FPR, which would not be possible in a blind test.

**Figure 5:**
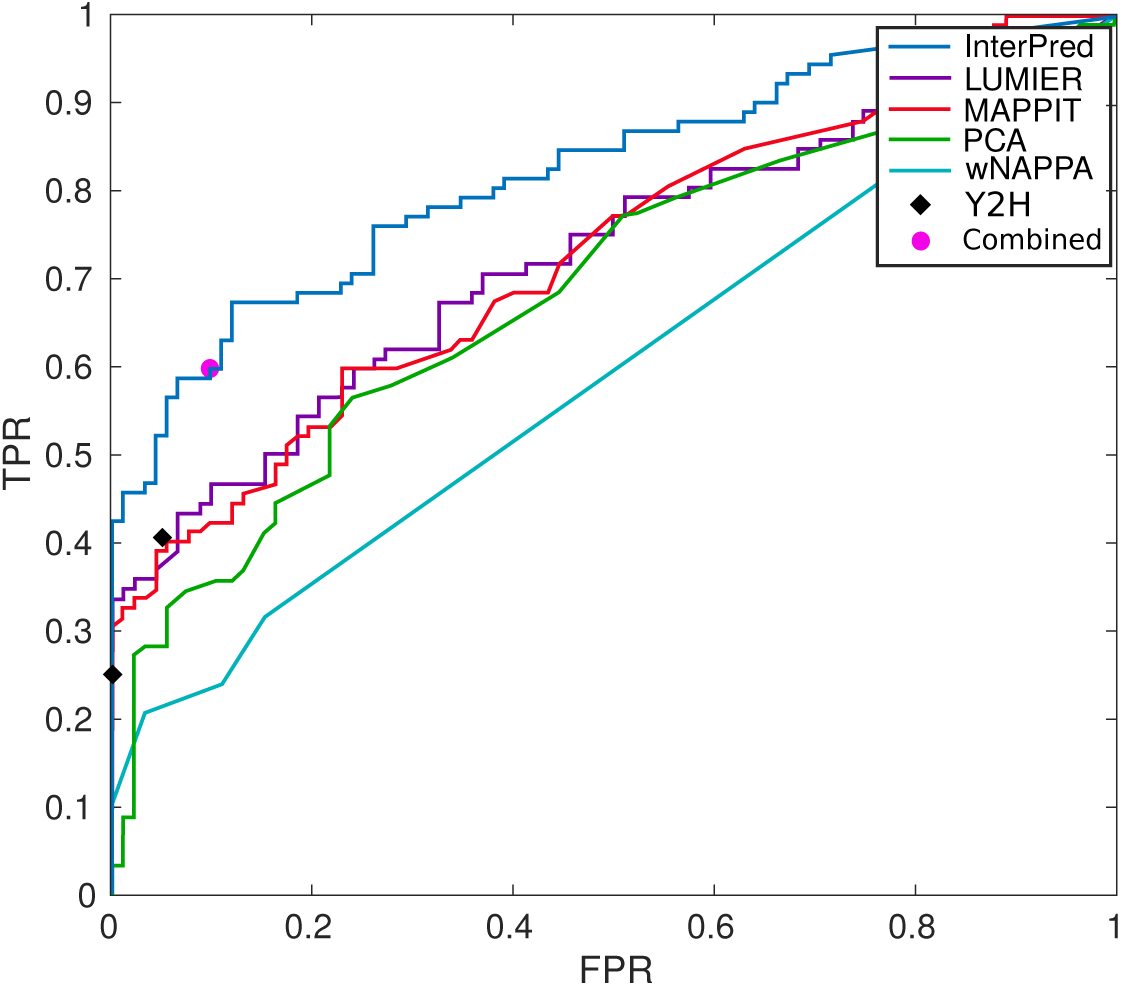
ROC curves comparing InterPred to a number of high-throughput methods on the Braun benchmark of 184 human protein pairs. The two black diamonds show the Y2H at two different bait and prey expression levels. The pink circle shows the performance if the experimental methods are combined in an optimal way.

### 3.4 Coarse modelling: Impact of using structure instead of sequence

A common technique to construct models of protein-protein interactions is to use template-based docking. It is conceptually similar to the InterPred pipeline, but with the important difference that only the sequence is used to find remote homologous structures. Here, we use the sequence to build a 3D model and search with the structure to find not only remote homologous but also similar structural interfaces.

To measure the added value of using structural information, we performed a “template reduction”, by removing template pairs at different thresholds from 90% to 20% sequence similarity, and finally templates that cannot be detected at all by PSI-BLAST (NA, E-value > 0.1). Figure 6 shows the percentage of targets from the BM4 set (see Methods) that could be correctly modelled, i.e. coarse interaction model with Interface Similarity score (IS-score) [48] ≥ 0.12, at different sequence similarity thresholds; by requiring that only one template (yellow) or both templates (blue) to be under the sequence similarity threshold. In reality the first case is probably most realistic, since to properly model an interaction both templates needs to be found. The results show that approximately a quarter (26%) of the targets could be modelled even when all templates found by PSI-BLAST are removed (“NA” bars). In the twilight zone, around 30% sequence similarity, InterPred can find structural templates for up to 40% of the targets in the benchmark. Thus, the added value of using structural information is the ability to model about 26%-40% of the proteins that would not be modelled otherwise.

**Figure 6.**
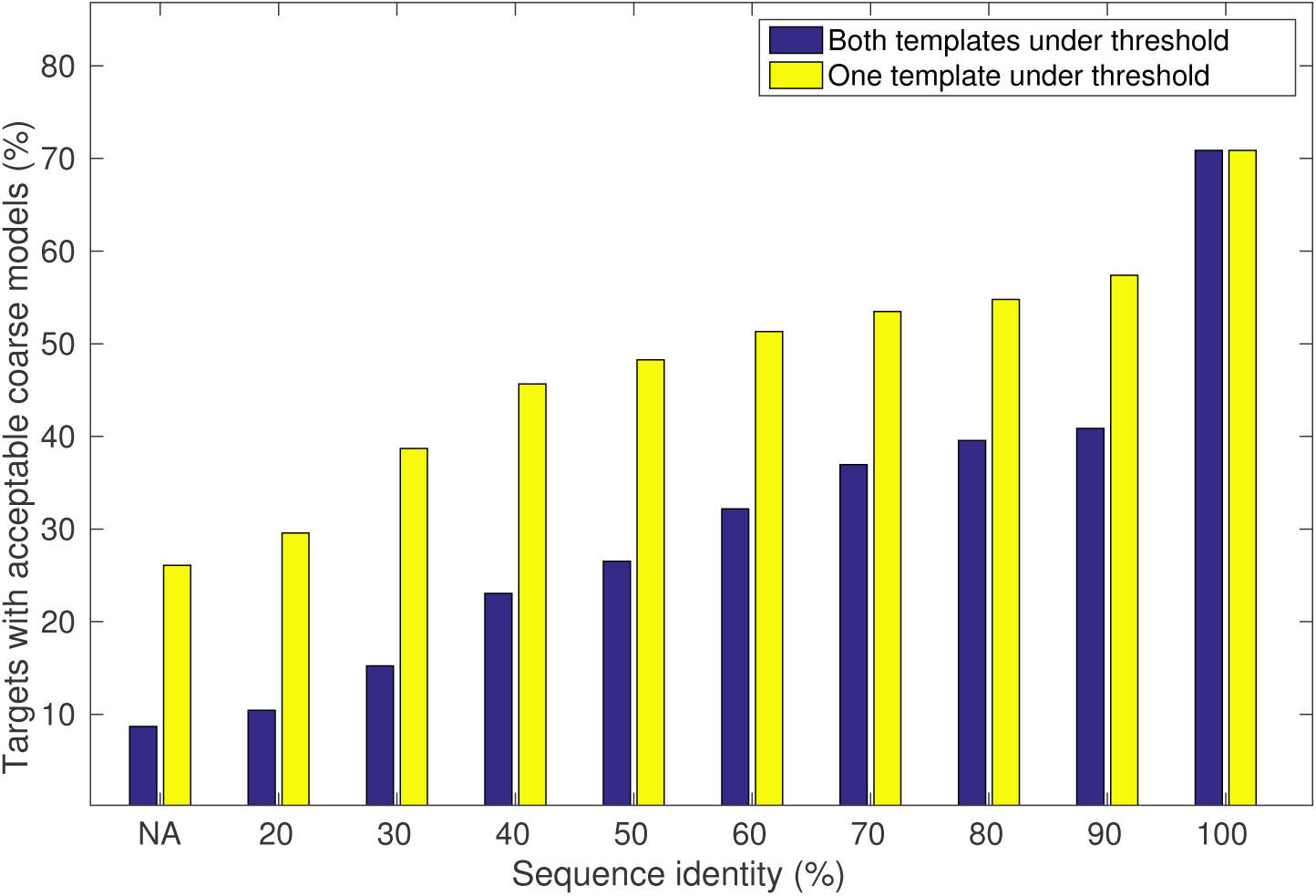
Percentage of acceptable coarse models built using templates with different sequence identity.

### 3.5 Coarse modelling: Evaluating the InterPred score

Next, we wanted to evaluate how well the InterPred score correlates with the effective quality of the coarse models that are produced when no obvious templates are available. To this end, we filtered out all templates with >30% sequence identity to any of the target pairs in BM4.

InterPred was used to construct coarse interaction models that were scored using the random forest. For each target, the top 10 coarse models by InterPred score were compared to the native complex using IS-score. In Figure 7 we show the percentage of targets that can be modelled with at least one acceptable model (IS-score ≥ 0.12) among the top 10 for different InterPred score thresholds. Each bin contains a subset of targets for which InterPred could find a model at a given InterPred score. To compare, we also used models generated with ZDOCK 3.0.2 using 15 degree sampling, downloaded from the ZDOCK webpage (see Methods).

**Figure 7.**
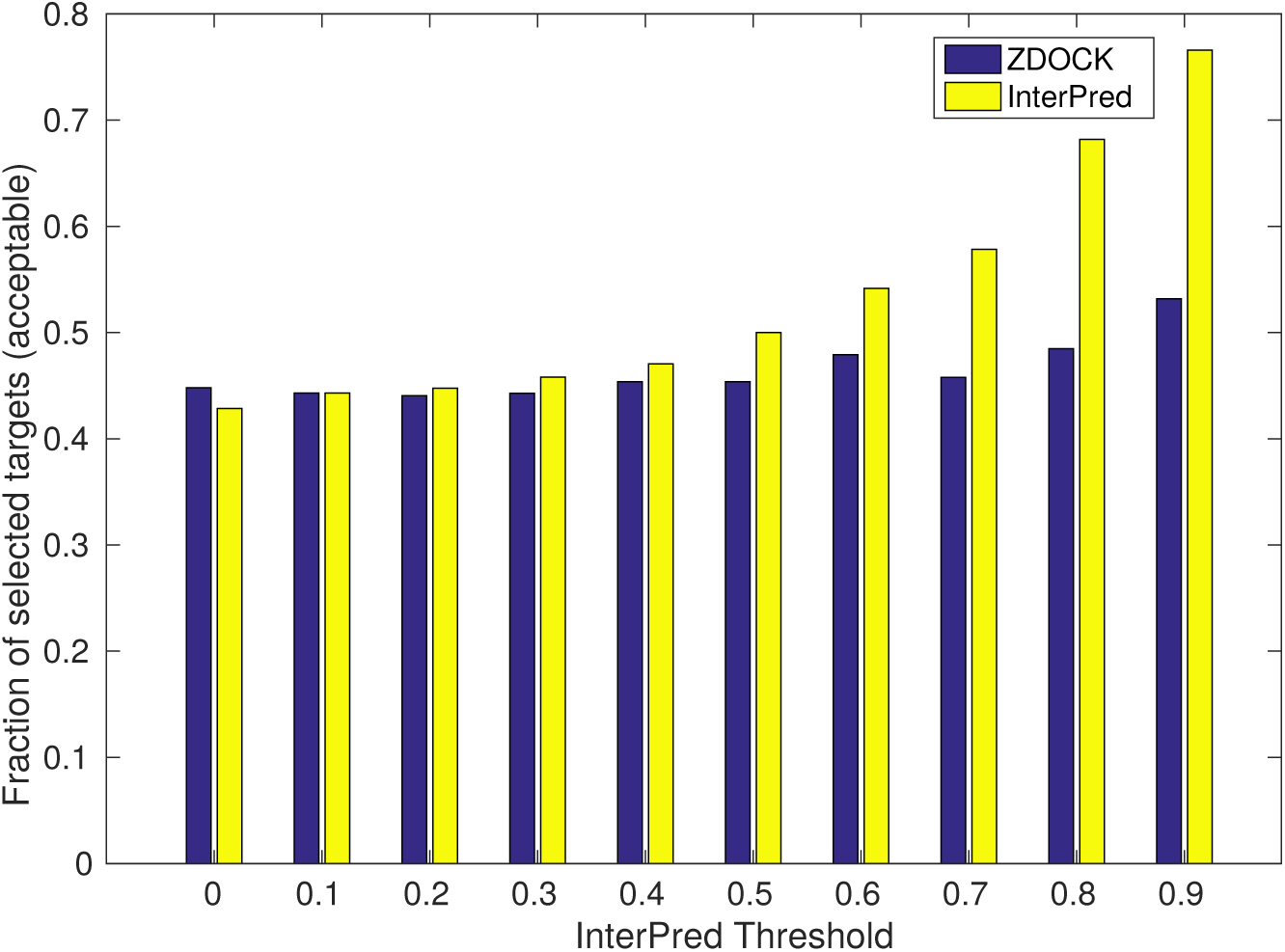
Fraction of correctly (acceptable or better) modelled targets from BM4 by InterPred (yellow) for different InterPred score excluding any template with sequence identity >30%. ZDOCK (blue) is included as a reference using the targets present in each bin. A target is considered correct if it has at least one acceptable model among the top 10.

ZDOCK is a template-free method for docking, it is still one of the most widely used rigid-body docking methods that are currently available for academic use. Moreover, template-free docking is usually the obvious choice when no obvious templates are available for a given complex such as in this case, where all templates with >30% sequence identity have been removed.

The fraction of acceptable models for InterPred increases from 45% for the lowest thresholds to around 80% for the highest. Thus, it is clear that the higher the InterPred score, the higher is the chance that the interaction model is correct.

If we compare InterPred and ZDOCK, the two methods perform similar up to InterPred score 0.4. But for InterPred score > 0.4 it progressively becomes more and more advantageous to use InterPred, as the fraction of correctly predicted targets peaks at 78% when at least one model with InterPred score ≥ 0.9 is available. The performance of ZDOCK remains roughly constant around 45% across all thresholds, meaning that the performance is stable across the targets.

### 3.6 Docking Refinement

In the final step of the InterPred pipeline the coarse interaction models are used as *starting points* for docking using the RosettaDock low and high-resolution protocol. To assess this step the 176 test cases in BM4 were used as input to InterPred. In this case, the inputs to the InterPred pipeline are the X-ray structures of the unbound targets as listed in the benchmark, thus there is no need to run the first homology modelling step of the pipeline. After the structural template search step, and removal of self-hits, the coarse models were ranked by InterPred score and the top 10 coarse models were selected for refinement.

Each selected coarse interaction model was used as a starting structure in a docking simulation using RosettaDock (see Methods). For each starting structure 10,000 decoys were generated (e.g. up to 100,000 decoys per test case if 10 coarse models are available) and their quality was evaluated using IS-score (see Methods).

We investigated several measures to rank the docked models (decoys), including the Rosetta energy score, interface score (Isc), IRMSD and RMSD between the docked model and the corresponding starting model. In Figure 8 we show a scatter plot with IS-score before and after docking, for models selected by Rosetta Score, IRMSD, and as comparison also the best docked model. In most cases the best docked model is better than the starting model. However, it is clear that neither the Rosetta energy score nor interface score (data not shown) is able to select these good models. In fact, the best measure we came up with is to select the models whose interface changes the least, i.e. the models with the lowest IRMSD to the starting structure.

**Figure 8:**
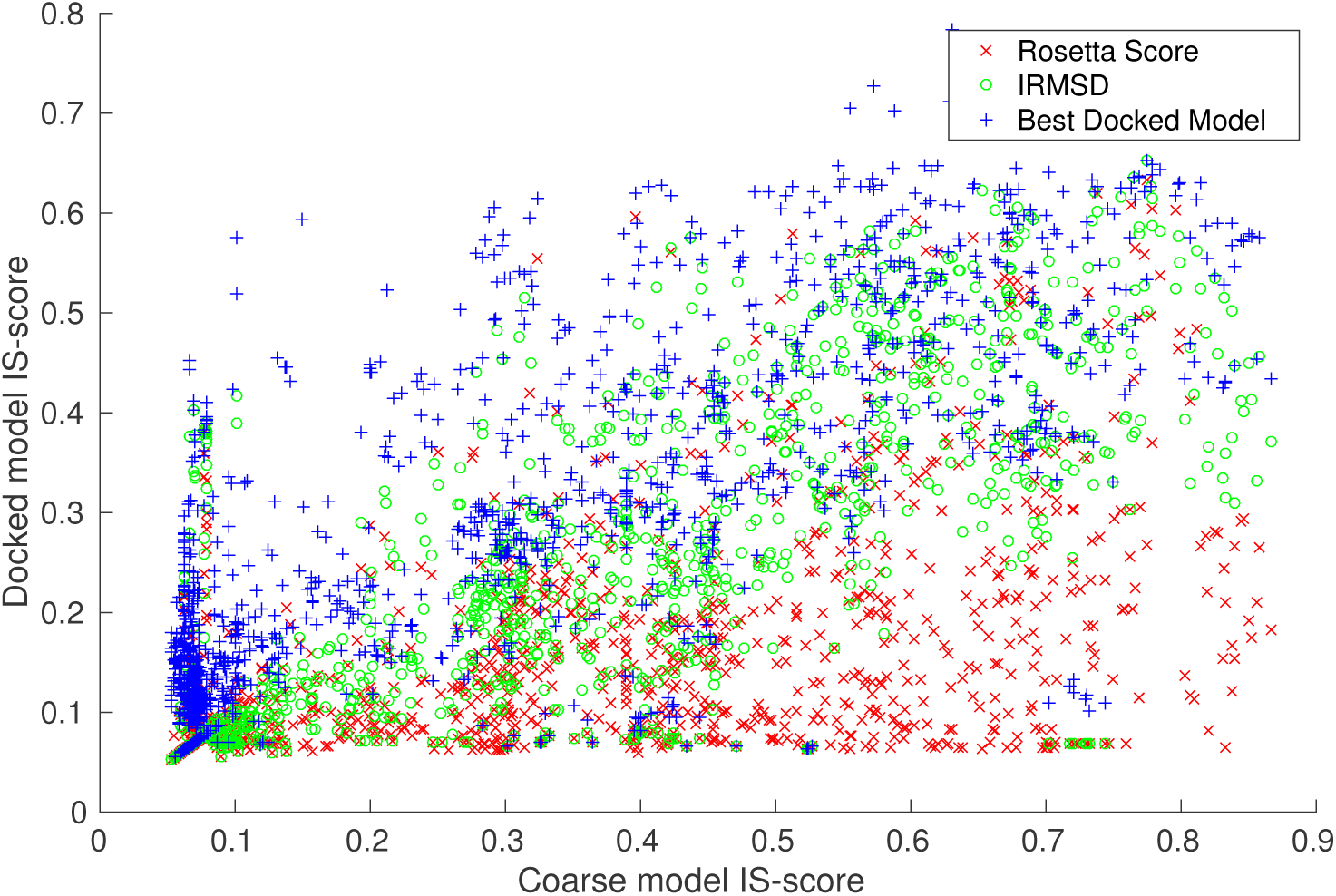
IS-scores before and after docking for the top 10 docked models ranked by Rosetta Score (red) or IRMSD (green), as reference the best IS-score (blue) model is also included.

The quality of the docked models was benchmarked against PRISM [41] and ZDOCK [50]. PRISM is most similar to InterPred in that it is also using structural templates (see Methods). ZDOCK is one of the leading *ab initio* methods using rigid-body docking. The results showing the percentage of test cases for which at least one Acceptable or Near-native model is produced among the top 1 to 1,000 models is shown in Figure 9.

**Figure 9.**
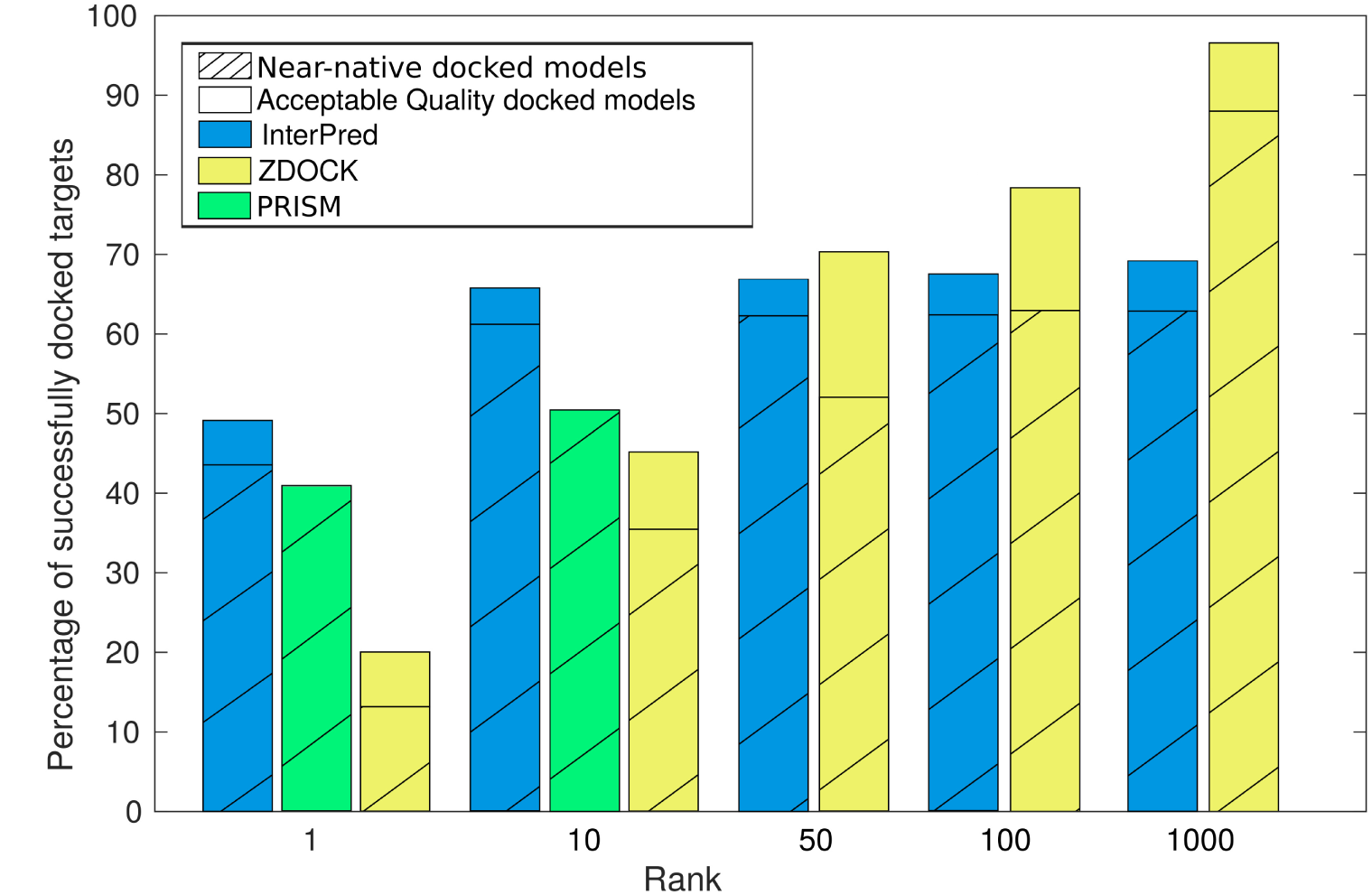
The percentage of successfully docked targets for InterPred (blue), PRISM (green) and ZDOCK (yellow). PRISM results is only available for rank 1 and 10, since it never returns more than 20 models per target.

PRISM never reported more than 20 models for any of the targets, thus the performance is only evaluated for the top 1 and top 10 ranked models. Compared to PRISM, InterPred performs considerably better having 85 out of 174 (49%) targets with an at least Acceptable model for top 1, compared to 72 (41%) Acceptable targets for PRISM. For top 10 the numbers are 113 (65%) for InterPred vs. 86 (50%) For PRISM. The difference is smaller when considering the Near-native quality models, as all Acceptable model by PRISM are also Near-native. For top 1 it is 75 (43%) vs. 72 (41%), for top 10 increasing to 106 (61%) vs. 86 (50%), for InterPred and PRISM, respectively.

The results for ZDOCK, being and *ab initio* method is obviously worse considering the top 1 and top 10 ranked models. However, starting from rank 50 ZDOCK performs better. The reason for this is that while the InterPred models are docked locally around a given starting point, ZDOCK samples the whole conformational space among its 3,600 predictions. This means that by considering lower ranked docked models you will almost always find a correct model. However, we would argue that it is not really realistic to consider more than the top 10 ranked models.

### 3.7 CAPRI37: Blind predictions

CAPRI (Critical Assessment of PRediction of Interactions) is a community-wide experiment on the comparative evaluation of protein-protein docking for structure prediction [42]. In 2016, the 37th round of CAPRI was held in collaboration with the 12th edition of CASP (Critical Assessment of protein Structure Prediction). Ten target multimeric complexes were released to the prediction community before the structure was solved experimentally, thus the predictions are completely blind predictions. A total of 17 different target interfaces were evaluated, since some of the target complexes included trimers, tetramers and octamers.

InterPred participated in CAPRI37 with a slightly modified version. Instead of running the first step of the pipeline (Homology Modelling), the server models of the target sequences were downloaded from the CASP webpage (http://predictioncenter.org) and evaluated by the Model Quality Assessment Program (MQAP) method Pcomb [57]. The top ranked models were then used in the structural template search.

The results for CAPRI37 are summarized in Table 2, following the CAPRI criteria that classify a docked model as either Incorrect, Acceptable, Medium, or High Quality, depending on the fraction of correctly predicted interfacial residues (FNat), RMSD of the model ligand to the native upon superposition of the receptors (LRMSD) and RMSD of the interfacial residues (IRMSD) [42].

**Table 2.**
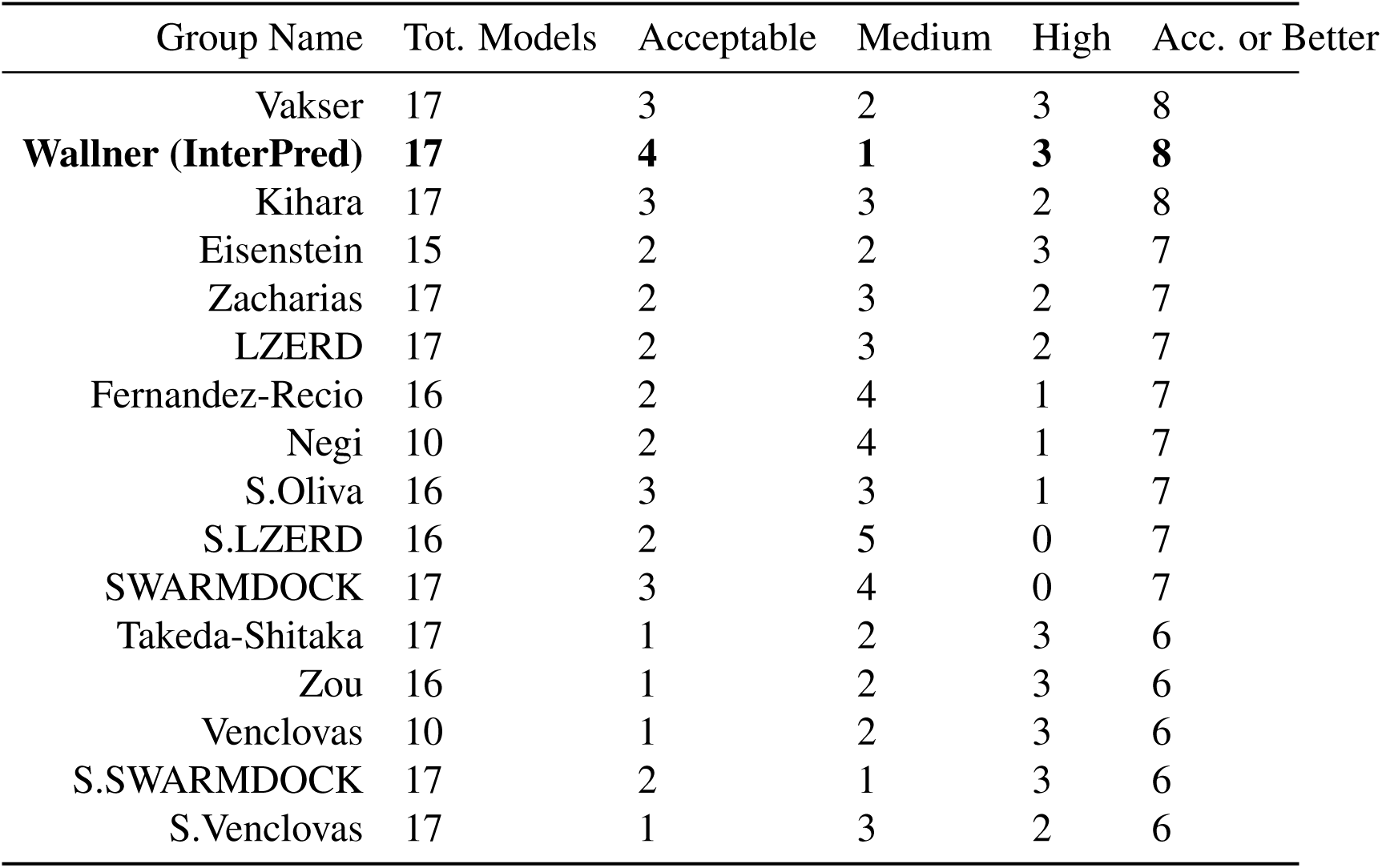
Table showing the top ranking groups in the 37th round of the CAPRI experiment, sorted by number of “Acceptable or Better” models.

| Group Name | Tot. Models | Acceptable | Medium | High | Acc. or Better |
| --- | --- | --- | --- | --- | --- |
| Vakser | 17 | 3 | 2 | 3 | 8 |
| **Wallner (InterPred)** | **17** | **4** | **1** | **3** | **8** |
| Kihara | 17 | 3 | 3 | 2 | 8 |
| Eisenstein | 15 | 2 | 2 | 3 | 7 |
| Zacharias | 17 | 2 | 3 | 2 | 7 |
| LZERD | 17 | 2 | 3 | 2 | 7 |
| Fernandez-Recio | 16 | 2 | 4 | 1 | 7 |
| Negi | 10 | 2 | 4 | 1 | 7 |
| S.Oliva | 16 | 3 | 3 | 1 | 7 |
| S.LZERD | 16 | 2 | 5 | 0 | 7 |
| SWARMDOCK | 17 | 3 | 4 | 0 | 7 |
| Takeda-Shitaka | 17 | 1 | 2 | 3 | 6 |
| Zou | 16 | 1 | 2 | 3 | 6 |
| Venclovas | 10 | 1 | 2 | 3 | 6 |
| S.SWARMDOCK | 17 | 2 | 1 | 3 | 6 |
| S.Venclovas | 17 | 1 | 3 | 2 | 6 |

As it is done in the CASP experiments, only the rank 1 model is considered for each target. InterPred (group name: Wallner) correctly predicted the most interactions, 8 “Acceptable or better“ models out of 17 target interfaces, along with groups of Kihara and Vakser.

### 3.8 Computational Cost

The two steps in the InterPred pipeline that are most computational expensive are the structural template search and the final docking step. Both these steps are trivial to parallelize, since they carry out many independent calculations. However to get some idea of the computational cost, we report the actual timings when run on a 16 core 2.2GHz with 32GB RAM Linux node using some typical size. Structural template search using a 400 residues target take on average 23 minutes (46 minutes for a target couple in total). The final docking step take around 180 minutes to generate 10,000 docking decoys for each coarse model. Thus, total time for a typical query would be 50+10×180 minutes or roughly 31h on a single compute node. On the other hand, as we have seen from Figure 9, there is not much of a difference between considering the top 10 or top 100 (or even 1000) docked decoys in terms of fraction of correctly predicted interactions. Thus, one might argue that producing only 1,000 decoys instead of 10,000 would generate approximately the same results while at the same time reducing the computational cost 10 times to around 3h.

## 4 Conclusions

We have presented InterPred, a tool that predicts if two proteins are interacting from their sequence and builds a full-atom model of the interaction. Starting from two protein sequences, 3D models are constructed and the structural neighborhoods of each 3D model are explored by comparing them against all possible structural domains in the PDB using structural alignment. Candidate coarse interaction models are constructed by superposition whenever structural neighborhoods coincide. These models are ranked using a novel random forest classifier that distinguish correct from incorrect interactions based on features calculated from the coarse interaction models. Finally, if the goal is to generate an all-atom description of the interaction, the top-ranked models are used as starting points in an all-atom docking procedure to generate the final docked conformation selected as the lowest IRMSD to the starting structure.

Our results show that the use of close and remote structural interaction templates represents a major improvement when comparing to methods where only the sequence (and/or sequence profiles) are used to predict interactions.

In addition, we also show that using InterPred to detect protein-protein interactions is even better than several experimental high-throughput methods, finding 25% more true interactions compared to the best high-throughput methods in low FPR region. In fact, only a consensus approach combining five high-throughput method in an optimal way is able to perform on par with InterPred.

The InterPred score is also a useful predictor of the success rate for modeling the protein-protein interaction in molecular detail starting from the coarse-grained interaction model. Molecular docking starting from interaction models with InterPred scores >0.5 produces acceptable quality models for 50% of the targets, while a score >0.9 produces acceptable quality models for almost 80% of the targets.

To test this further InterPred was used to generate full-atom docked models on the Docking Benchmark 4.0, and showed that it performs better than other state of the art predictors based on structural alignments, (PRISM), with an improvement of almost 20% for the first ranked decoy, and the best available docking pipeline (ZDOCK 3.0.2) for the top ranked docking decoys with an improvement of around 150% for the first ranked decoy. Overall, 45-50% target interactions were correctly modelled.

The fact that InterPred performs on par on better than state of the art was also confirmed in the latest CAPRI37 experiment, where InterPred could correctly predict, roughly half of the target interfaces, ranking amongst the top predictors in the community.

InterPred is available as a standalone download from http://wallnerlab.org/InterPred/, and should be useful for anyone working with protein-protein interactions.

## 5 Acknowledgement

This work was supported by Swedish Research Council grants 2012-5270, 2016-05369, The Swedish e-Science Research Center, and the Foundation Blanceflor Boncompagni Ludovisi, née Bildt. The computations were performed on resources provided by the Swedish National Infrastructure for Computing (SNIC) at the National Supercomputer Centre (NSC) in Linköping.

## References

1. Wang, P. I. and Marcotte, E. M. It’s the machine that matters: Predicting gene function and phenotype from protein networks. Journal of proteomics 73(11):2277–2289, October, 2010.

2. Shoemaker, B. A. and Panchenko, A. R. Deciphering protein-protein interactions. Part I. Experimental techniques and databases. PLoS computational biology 3(3):e42, March, 2007.

3. Parrish, J. R., Gulyas, K. D., and Finley, R. L. Yeast two-hybrid contributions to interactome mapping. Current opinion in biotechnology 17(4):387–393, August, 2006.

4. Vasilescu, J. and Figeys, D. Mapping protein-protein interactions by mass spectrometry. Current opinion in biotechnology 17(4):394–399, August, 2006.

5. Braun, P. Interactome mapping for analysis of complex phenotypes: insights from benchmarking binary interaction assays. Proteomics 12(10):1499–1518, May, 2012.

6. Roux, K. J., Kim, D. I., and Burke, B. BioID: a screen for protein-protein interactions. Current protocols in protein science / editorial board, John E., Coligan… [et al.] 74:Unit 19.23, 2013.

7. Lu, L., Lu, H., and Skolnick, J. MULTIPROSPECTOR: an algorithm for the prediction of protein-protein interactions by multimeric threading. Proteins: Structure, Function, and Bioinformatics 49(3):350–364, November, 2002.

8. You, Z.-H., Lei, Y.-K., Zhu, L., Xia, J., and Wang, B. Prediction of protein-protein interactions from amino acid sequences with ensemble extreme learning machines and principal component analysis. BMC bioinformatics 14 Suppl(8):S10, 2013.

9. Hamp, T. and Rost, B. Evolutionary profiles improve protein-protein interaction prediction from sequence. Bioinformatics 31(12):1945–1950, 2015.

10. Pazos, F. and Valencia, A. Similarity of phylogenetic trees as indicator of protein-protein interaction. Protein Eng 14(9):609–614, September, 2001.

11. Overbeek, R., Fonstein, M., D’Souza, M., Pusch, G. D., and Maltsev, N. The use of gene clusters to infer functional coupling. Proceedings of the National Academy of Sciences of the United States of America 96(6):e2896–2901, March, 1999.

12. Pellegrini, M., Marcotte, E. M., Thompson, M. J., Eisenberg, D., and Yeates, T. O. Assigning protein functions by comparative genome analysis: protein phylogenetic profiles. Proceedings of the National Academy of Sciences of the United States of America 96(8):4285–4288, April, 1999.

13. Zhang, Q. C., Petrey, D., Garzón, J. I., Deng, L., and Honig, B. PrePPI: a structure-informed database of protein-protein interactions. Nucleic acids research 41(Database issue):D828–33, January, 2013.

14. Chen, H. and Skolnick, J. M-TASSER: an algorithm for protein quaternary structure prediction. Biophys J 94(3):918–928, Feb, 2008.

15. Mukherjee, S. and Zhang, Y. Protein-protein complex structure predictions by multimeric threading and template recombination. Structure 19(7):955–966, Jul, 2011.

16. Guerler, A., Govindarajoo, B., and Zhang, Y. Mapping monomeric threading to protein-protein structure prediction. J Chem Inf Model 53(3):717–725, Mar, 2013.

17. Davis, F. P., Braberg, H., Shen, M.-Y., Pieper, U., Sali, A., and Madhusudhan, M. S. Protein complex compositions predicted by structural similarity. Nucleic acids research 34(10):2943–2952, 2006.

18. Kundrotas, P. J. and Vakser, I. A. Global and local structural similarity in protein-protein complexes: Implications for template-based docking. Proteins: Structure, Function, and Bioinformatics 81(12):2137–2142, 2013.

19. Gao, M. and Skolnick, J. ialign: a method for the structural comparison of protein-protein interfaces. Bioinformatics 26(18):2259–2265, 2010.

20. Sinha, R., Kundrotas, P. J., and Vakser, I. A. Docking by structural similarity at protein-protein interfaces. Proteins: Structure, Function, and Bioinformatics 78(15):3235–3241, 2010.

21. Tuncbag, N., Gursoy, A., Nussinov, R., and Keskin, O. Predicting protein-protein interactions on a proteome scale by matching evolutionary and structural similarities at interfaces using PRISM. Nat Protoc 6(9):1341–1354, August, 2011.

22. Baspinar, A., Cukuroglu, E., Nussinov, R., Keskin, O., and Gursoy, A. Prism: a web server and repository for prediction of protein-protein interactions and modeling their 3d complexes. Nucleic acids research 42(W1):W285–W289, 2014.

23. Dominguez, C., Boelens, R., and Bonvin, A. M. J. J. HADDOCK: A Protein-Protein Docking Approach Based on Biochemical or Biophysical Information. Journal of the American Chemical Society 125(7):1731–1737, February, 2003.

24. Kozakov, D., Brenke, R., Comeau, S. R., and Vajda, S. Piper: an fft-based protein docking program with pairwise potentials. Proteins: Structure, Function, and Bioinformatics 65(2):392–406, 2006.

25. Mintseris, J., Pierce, B., Wiehe, K., Anderson, R., Chen, R., and Weng, Z. Integrating statistical pair potentials into protein complex prediction. Proteins: Structure, Function, and Bioinformatics 69(3):511–520, 2007.

26. Pierce, B. G., Wiehe, K., Hwang, H., Kim, B.-H., Vreven, T., and Weng, Z. Zdock server: interactive docking prediction of protein-protein complexes and symmetric multimers. Bioinformatics 30(12):1771–1773, 2014.

27. Moreira, I. S., Fernandes, P. A., and Ramos, M. J. Protein-protein docking dealing with the unknown. J Comput Chem 31(2):317–342, Jan, 2010.

28. Huang, S.-Y. Exploring the potential of global protein-protein docking: an overview and critical assessment of current programs for automatic ab initio docking. Drug Discovery Today 20(8):969–977, 2015.

29. Tuncbag, N., Gursoy, A., Guney, E., Nussinov, R., and Keskin, O. Architectures and functional coverage of protein-protein interfaces. Journal of Molecular Biology 381(3):785–802, September, 2008.

30. Zhang, Q. C., Petrey, D., Norel, R., and Honig, B. H. Protein interface conservation across structure space. Proceedings of the National Academy of Sciences of the United States of America 107(24):10896–10901, June, 2010.

31. Gao, M. and Skolnick, J. Structural space of protein-protein interfaces is degenerate, close to complete, and highly connected. Proceedings of the National Academy of Sciences of the United States of America 107(52):22517–22522, December, 2010.

32. Zhang, Q. C., Petrey, D., Deng, L., Qiang, L., Shi, Y., Thu, C. A., Bisikirska, B., Lefebvre, C., Accili, D., Hunter, T., Maniatis, T., Califano, A., and Honig, B. Structure-based prediction of protein-protein interactions on a genomewide scale. Nature 490(7421):556–560, October, 2012.

33. Sinha, R., Kundrotas, P. J., and Vakser, I. A. Protein docking by the interface structure similarity: how much structure is needed? PloS one 7(2):e31349, 2012.

34. Tuncbag, N., Keskin, O., Nussinov, R., and Gursoy, A. Fast and accurate modeling of protein-protein interactions by combining template-interface-based docking with flexible refinement. Proteins: Structure, Function, and Bioinformatics 80(4):1239–1249, 2012.

35. Szilagyi, A. and Zhang, Y. Template-based structure modeling of protein-protein interactions. Curr Opin Struct Biol 24:10–23, Feb, 2014.

36. Berman, H. M., Westbrook, J., Feng, Z., Gilliland, G., Bhat, T. N., Weissig, H., Shindyalov, I. N., and Bourne, P. E. The Protein Data Bank. Nucleic acids research 28(1):235–242, January, 2000.

37. Remmert, M., Biegert, A., Hauser, A., and Söding, J. HHblits: lightning-fast iterative protein sequence searching by HMM-HMM alignment. Nature methods 9(2):173–175, February, 2012.

38. Webb, B. and Sali, A. Comparative Protein Structure Modeling Using MODELLER. Current protocols in bioinformatics / editoral board, Andreas D. Baxevanis… [et al] 47:5.6.1–5.6.32, 2014.

39. Zhang, Y. and Skolnick, J. TM-align: a protein structure alignment algorithm based on the TM-score. Nucleic acids research 33(7):2302–2309, 2005.

40. Anishchenko, I., Kundrotas, P. J., and Vakser, I. A. Modeling complexes of modeled proteins. Proteins: Structure, Function, and Bioinformatics 85:470–478, 2017.

41. Ogmen, U., Keskin, O., Aytuna, A. S., Nussinov, R., and Gursoy, A. Prism: protein interactions by structural matching. Nucleic acids research 33(suppl 2):W331–W336, 2005.

42. Lensink, M. F., Méndez, R., and Wodak, S. J. Docking and scoring protein complexes: CAPRI 3rd Edition. Proteins: Structure, Function, and Bioinformatics 69(4):704–718, December, 2007.

43. Gray, J. J., Moughon, S., Wang, C., Schueler-Furman, O., Kuhlman, B., Rohl, C. A., and Baker, D. Protein-protein docking with simultaneous optimization of rigid-body displacement and side-chain conformations. Journal of Molecular Biology 331(1):281–299, August, 2003.

44. Gene Ontology Consortium. Gene Ontology Consortium: going forward. Nucleic acids research 43(Database issue):D1049–56, January, 2015.

45. Altschul, S. F., Madden, T. L., Schaffer, A. A., Zhang, J., Zhang, Z., Miller, W., and Lipman, D. J. Gapped BLAST and PSI-BLAST: a new generation of protein database search programs. Nucleic acids research 25(17):3389–3402, 1997.

46. Braun, P., Tasan, M., Dreze, M., Barrios-Rodiles, M., Lemmens, I., Yu, H., Sahalie, J. M., Murray, R. R., Roncari, L., De Smet, A.-S., et al. An experimentally derived confidence score for binary protein-protein interactions. Nature methods 6(1):91–97, 2009.

47. Hwang, H., Vreven, T., Janin, J., and Weng, Z. Protein-protein docking benchmark version 4.0. Proteins 78(15):3111–3114, Nov, 2010.

48. Gao, M. and Skolnick, J. New benchmark metrics for protein-protein docking methods. Proteins: Structure, Function, and Bioinformatics 79(5):1623–1634, 2011.

49. Tuncbag, N., Gursoy, A., Nussinov, R., and Keskin, O. Predicting protein-protein interactions on a proteome scale by matching evolutionary and structural similarities at interfaces using prism. Nature protocols 6(9):1341–1354, 2011.

50. Pierce, B. G., Hourai, Y., and Weng, Z. Accelerating protein docking in ZDOCK using an advanced 3D convolution library. PloS one 6(9):e24657, 2011.

51. ZDOCK Decoy Sets Webpage. https://zlab.umassmed.edu/zdock/decoys.shtml.

52. Walhout, A. J. and Vidal, M. High-throughput yeast two-hybrid assays for large-scale protein interaction mapping. Methods 24(3):297–306, 2001.

53. Eyckerman, S., Verhee, A., Van der Heyden, J., Lemmens, I., Van Ostade, X., Vandekerckhove, J., and Tavernier, J. Design and application of a cytokine-receptor-based interaction trap. Nature Cell Biology 3(12):1114–1119, 2001.

54. Barrios-Rodiles, M., Brown, K. R., Ozdamar, B., Bose, R., Liu, Z., Donovan, R. S., Shinjo, F., Liu, Y., Dembowy, J., Taylor, I. W., et al. High-throughput mapping of a dynamic signaling network in mammalian cells. Science 307(5715):1621–1625, 2005.

55. Nyfeler, B., Michnick, S. W., and Hauri, H.-P. Capturing protein interactions in the secretory pathway of living cells. Proceedings of the National Academy of Sciences of the United States of America 102(18):6350–6355, 2005.

56. Ramachandran, N., Raphael, J. V., Hainsworth, E., Demirkan, G., Fuentes, M. G., Rolfs, A., Hu, Y., and LaBaer, J. Next-generation high-density self-assembling functional protein arrays. Nature methods 5(6):535–538, 2008.

57. Wallner, B. and Elofsson, A. Prediction of global and local model quality in casp7 using pcons and proq. Proteins: Structure, Function, and Bioinformatics 69(S8):184–193, 2007.

